# Maternal immune activation in rats is associated with a lower number of dopamine receptor 3-expressing granulocytes in the male offspring with no alterations in cocaine reward, resistance to extinction or cue-induced reinstatement

**DOI:** 10.1101/796425

**Authors:** Raquel Santos-Toscano, Marcos Ucha, Érika Borcel, Emilio Ambrosio, Alejandro Higuera-Matas

**Author notes:** School of Medicine. University of Central Lancashire, Preston, UK. These authors contributed equally to this work (Raquel Santos-Toscano, Marcos Ucha). Corresponding authors at: Departamento de Psicobiología, Facultad de Psicología, UNED, C/Juan del Rosal 10, Madrid, Spain. E-mail addresses (E Ambrosio) & (A. Higuera-Matas).

## Abstract

There is evidence for increased rates of drug use among schizophrenic patients. However, the causality in this relationship remains unclear. In addition, biomarkers of schizophrenia are vital, given the heterogeneous nature of the disorder that can lead to difficulties in the early diagnosis. In the present work, we use a maternal immune activation model to experimentally test whether animals at high risk of developing a schizophrenia-like condition are more prone to acquire cocaine self-administration, show enhanced sensitivity to the reinforcing actions of cocaine or if they are resistant to extinction or vulnerable to relapse. Pregnant rats were injected with lipopolysaccharide (LPS) (2 mg/kg s.c.) or saline every other day during pregnancy, and the offspring was tested for sensorimotor gating (prepulse inhibition –PPI-). After this test, one group of rats was submitted to cocaine self-administration (0.5 mg/kg) under fixed and progressive ratio schedules, dose-response testing, extinction and cue-induced drug-seeking. Another group was sacrificed to study potential biomarkers in the immune blood cells by flow cytometry. While rats born to LPS-treated mothers showed impaired PPI, there were no differences in cocaine self-administration acquisition, responsiveness to dose shifts, extinction or cue-induced reinstatement. Finally, there were fewer DRD3^+^ granulocytes in the LPS-offspring and an exciting trend for CNR2^+^ lymphocytes to be more abundant in LPS-exposed rats. Our results indicate that the higher prevalence of cocaine abuse among people with schizophrenia is not due to a pre-existing pathology and suggest that DRD3^+^ granulocytes and possibly CNR2^+^ lymphocytes could be potential biomarkers of schizophrenia.

## Introduction

Schizophrenia is a complex mental disorder that affects 23 million worldwide (World Health Organization, 2018). This condition is characterized by positive, negative, cognitive and mood symptoms affecting perception, language, thought and overt behaviour. Due to the high heterogeneity of this wide cluster of symptoms among the patients, the diagnosis is a challenge for the clinicians (Gaebel and Zielasek, 2015).

There is a high prevalence of substance use disorders (SUDs) among people with schizophrenia, which tend to worsen as the disease progresses. Although the drugs more frequently abused by this population are nicotine and caffeine, others such as alcohol, cannabis and cocaine are not uncommon. Comorbidity with cocaine use disorder has not been sufficiently studied as compared to other drugs. However, the prevalence of cocaine use disorder has been found to range between 15% and 50% in schizophrenia patients (Thoma and Daum, 2013).

There are just a few studies that have analysed cocaine self-administration in animal models of schizophrenia. Chambers and Self, using the neonatal ventral hippocampal lesion (NVHL) observed a more robust acquisition of cocaine self-administration, resistance to extinction and high rate of relapse as compared with sham-operated animals (Chambers and Self, 2002). In the same vein, NVHL rats exhibited an increase in cue-induced reinstatement compared to the control group (Karlsson et al., 2013). Maternal immune activation (MIA) models recapitulate several aspects of schizophrenia and are rooted in the fact that prenatal infections are a predisposition factor to develop the condition (Brown and Meyer, 2018). These MIA models offer an excellent means to examine the causal relationships that exist between schizophrenia and the increased prevalence of addictive disorders in these patients.

Contrary to our expectations, we observed no increased cocaine self-administration in the offspring of rats exposed to LPS during pregnancy (Santos-Toscano et al., 2016). However, in this previous study, we did not test for other aspects cocaine-related behaviours that are relevant to addiction such as sensitivity to the rewarding actions of the drug in a dose-response setting, motivation to obtain the drug, resistance to extinction or vulnerability to cue-induced relapse. The main goal of the experiments here reported is to examine these behaviours.

Due to the strong relationship between the dopaminergic system and schizophrenia, certain components of this neurotransmission pathway have been postulated as biomarkers of the disorder. In this regard, positive symptoms correlate positively with the levels of dopamine receptor 2 (D2R) mRNA in lymphocytes (Liu et al., 2013). Also, the severity of symptoms correlates with DRD3 mRNA levels in lymphocytes (Kwak et al., 2001). The mRNA expression level of DRD3 in T lymphocytes was also significantly higher in patients with schizophrenia than in controls (Cui et al., 2015). In another study, DRD2 mRNA levels in lymphocytes were increased in schizophrenic patients, while DRD4 mRNA levels in CD4^+^ T cells were decreased (Boneberg et al., 2006).

There is growing evidence that supports the relationship between the endocannabinoid system and schizophrenia (Bossong et al., 2014; Fakhoury, 2017; Gupta et al., 2014; Ibarra-Lecue et al., 2018; Saito et al., 2013). Moreover, the activation of this system produces the release of circulating cytokines and different neurotransmitters, including dopamine (Foster et al., 2016; Kucerova et al., 2014). Therefore, the endocannabinoid system could also act as a possible biomarker for this disorder. As regards this, a decrease in the cannabinoid receptor 2 (CNR2) mRNA in the peripheral blood mononuclear cells (PBMC) has been found both in the first outbreaks and in chronic patients (Bioque et al., 2013; Ferretjans et al., 2014). In another study, patients with schizophrenia showed an increase in CNR1 expression in CD4+ T-cells, while the CNR1 and CNR2 expression in B-cells was decreased (Campos-carli et al., 2017). Regarding schizophrenic symptoms, the levels of CNR1 and CNR2 mRNA in PBMCs were significantly higher in schizophrenic patients compared to controls and these levels correlated positively with positive and negative schizophrenia symptoms and negatively with cognitive functioning (Chase et al., 2016).

In the light of these data, we decided to examine the relative levels of DRD3^+^ and CNR2^+^ immune cells in the MIA model as a proof of concept of their utility as potential biomarkers of schizophrenia.

## Materials and methods

### 1. Animals

The experiments described here were performed on the 8–10-week-old male offspring of Lewis rats obtained from Charles River (France). All animals were housed in transparent Plexiglas cages and kept in a temperature and humidity-controlled environment (21±1 °C/50–60%), with artificial lighting (12 h/12 h light/dark cycle, lights on at 8 am), and ad libitum access to food (commercial diet for rodents A04: Panlab, Barcelona, Spain) and tap water.

### 2. Experimental design

Female rats were mated with males (2 females per male) one week after arrival to the animal facility. Vaginal smears were taken daily from breeder females and pregnancy was determined by the presence of sperm in the vaginal smear (=day 0 of pregnancy). LPS (Lipopolysaccharide from Escherichia coli, Sigma 129K 4089) was subcutaneously (s.c.) administered to the pregnant rats (LPS n= 7) at a dose of 2 mg/kg on alternate days during pregnancy (Romero et al., 2010). LPS was dissolved in saline solution (0.9% Sodium chloride (NaCl) in water) and administered in a volume of 1.5 ml/kg. The control group (SAL n= 7) consisted on pregnant rats submitted to the same treatment schedule with saline instead of LPS.

The litters were culled so that there were no more than 12 pups per dam. Only the male offspring were used in these experiments. Litters were left undisturbed until postnatal day (PD) 21 when they were weaned in groups of 2–3. All rats in a given cage were from the same litter and therefore the same treatment.

Rats were subjected to PPI test on PD 120 (SAL n= 37/LPS n= 22) and on PD 130 were divided into two groups: one of them was used for cocaine self-administration experiments (SAL n= 9/LPS n= 16), and the other was sacrificed for flow cytometry determinations (SAL n= 12/LPS n= 9). All behavioural tests were carried out during the light cycle, between 9 am and 5 pm.

### 3. Bodyweight

Maternal body weight was measured daily before each injection.

### 4. Prepulse inhibition of the acoustic startle response

PPI of the acoustic startle was measured in a non-restrictive Plexiglas cage (28 × 15 × 17 cm) that encloses the sensor’s platform (Cibertec, Madrid). If the animal moves up or down, a transient force is developed on the platform and measured at its peak, being equivalent in amplitude to the startle response (Romero et al., 2007).

Rats were placed in the startle chamber and each test session started with a 5-min acclimation period consisting of background white noise at 70 decibels (dB) (which continued throughout the session) to then be subjected to 52 startle trials under three conditions: (1) a main stimulus consisting on a 115-dB, 40-ms noise burst presented alone; (2) this same burst preceded by a 20-ms duration prepulse either 30 or 120 ms before the main stimulus and of an intensity of either 8 or 16 dB above background noise (30_8, 30_16, 120_8, 120_16 trial types); and (3) no-stimulus trials.

Five main stimuli (pulse alone) trials were presented at the beginning and at the end of each session to stabilise startle response and to calculate the habituation of such response but were not used for the calculation of PPI. In between these blocks, there were 42 mixed trials presented in a pseudorandom order: 12 pulse alone trials; 24 prepulse trials (six of each type); and six no-stimulus trials. The duration of the sessions was of around 15 min.

Prepulse inhibition is expressed as % PPI, and it was calculated using the following formula: 1−[startle amplitude on prepulse+pulse trial / mean startle amplitude on pulse alone trials]) × 100.

### 5. Cocaine self-administration

Subjects were surgically prepared with an intravenous catheter (0.064 i.d.) implanted in the right jugular vein approximately at the level of the atrium. The catheter was passed subcutaneously and exited in the mid scapular region. The animals were given at least seven days to recover after surgery. To prevent infections and to maintain catheter patency, the catheters were flushed daily with 0.5 ml of a solution of antibiotic (gentamicin, 40 mg/ml) dissolved in heparinised saline. Twelve operant chambers (Coulburn Instruments, Allentown, PA, USA) were used for cocaine self-administration studies equipped with two levers designed to register the animal responses. A microliter injection pump (Harvard 22) was used to deliver i.v. cocaine injections (0.5 mg/kg, for 4 seconds) when the active lever was pressured (inactive lever presses were recorded but had no programmed consequences). A light that was situated over the lever acted as a discriminative visual cue and was illuminated for 3 seconds whenever the active lever was pressed.

Self-administration sessions were carried out in 2 h daily sessions seven days a week for approximately seven weeks. The protocol was divided into several phases: 23 days of acquisition at 0.5 mg/kg dose under a fixed-ratio 1 schedule of reinforcement, a dose-response curve (one day per each dose of 1.5/1/0.25/0.125 mg/kg), three stabilization sessions with a 0.5 mg/kg dose, 6 days of progressive ratio schedule (Richardson and Roberts, 1996), another three stabilization sessions, 6 extinction sessions (where the drug was substituted with saline and the visual cue was not present) and cue-induced reinstatement (where the cue was activated, but the drug was not available).

### 6. Flow Cytometry

#### 6.1 Blood cell collection and red cells lysis

On PD 130, a different batch of animals was decapitated, and the peripheral blood was collected in EDTA tubes. The blood was centrifuged at 500g for 5min at 4°C and the plasma was separated. 1.5 ml of blood was then transferred to a new tube and 10 ml of 1X lysis buffer (Biolegend) was added and allowed to react with the sample for 23 min. The samples were then centrifuged again at 500g for 5 min (4°C). Three washes with phosphate buffer saline (PBS) were performed and after the last wash, the pellet was resuspended in 400 μl of complete RPMI medium (Sigma-Aldrich Co). The entire procedure was carried out at 4°C.

#### 6.2 Cell labelling

We used flow cytometry to study the populations of monocytes, granulocytes, and T and B lymphocytes, as well as the surface expression dopamine 3 (DRD3), dopamine 5 (DRD5) and cannabinoid 2 (CNR2) receptors. To this end, a sample volume containing 1×10^6^ cells was incubated with the primary antibody corresponding to each receptor diluted in PBS and 10% foetal bovine serum (Table 1) for 30 min. Two washes were then performed with PBS and incubated with a cocktail of antibodies labelled with different fluorochromes (Table 1) for another 30 min. After washing with PBS, the cells were fixed with FACSlysing solution (BD Biosciences) and analysed on a flow cytometer FACSCalibur (Becton Dickinson).

**Table 1.** Main characteristics and concentration of the antibodies used dor flow cytometry.

| Antigen | Target cell | Fluorophore | Host Species | Isotype | Concentration | Provider |
| --- | --- | --- | --- | --- | --- | --- |
| CD3 (rat) | T Lymphocytes | FITC | Mouse | IgG | 2.5µg/ml | Becton Dickinson |
| CD11b/c (rat) | Monocytes and granulocytes | PE | Mouse | IgG | 2.5µg/ml | Becton Dickinson |
| CD45RA (rat) | B Lymphocytes | PE-Cy™5 | Mouse | IgG | 1.25µg/ml | Becton Dickinson |
| CB2 Receptor |  |  | Rabbit | IgG | 10µg/ml | Thermo Scientific |
| D3 receptor |  |  | Rabbit | IgG | 15µg/ml | Abcam |
| D5 receptor |  |  | Rabbit | IgG | 15µg/ml | Abcam |
| IgG (rabbit) |  | Alexa Fluor 647 | Goat | IgG | 30µg/ml | Life Technologies |

For each sample, three tubes were tested, one for evaluating DRD3, DRD5 and CNR2 receptors (corresponding primary antibody: Table 1), and another control tube sample but no primary antibody to which the labelled secondary antibody (Table 1) could bind. The three antibodies that identify the cell populations were added to the control tubes in order to correct the potential signal due to the non-specific binding of the secondary antibody (Secondary antibody control).

### 7. Data Analysis

All data were analysed using the IBM SPSS statistical software package (version 24 for Windows). Outliers were identified by SPSS using the interquartile range criterion and a value of p ≤ 0.05 was considered to represent a statistically significant difference. Square roots transformations were applied when appropriate to correct the skewness in the distribution of the data and the lack of homogeneity of variances.

The increment in body weight was calculated for each day of injection, taking the weight of the previous injection day as a reference. Bodyweight increment was analysed by ANOVA (fixed between-group factor, Treatment [SAL or LPS]; fixed within-subject repeated measures factors, Injection Day).

PPI test was analysed with Student’s t-test for independent samples. Each PPI parameter was considered as a dependent variable and Treatment as an independent variable.

All phases in the cocaine self-administration study were analysed with ANOVA (fixed between-group factor, Treatment [SAL or LPS]; fixed within-subject repeated measures factors, Daily cocaine self-administration sessions). Cue-induced reinstatement was analysed with a Student’s t-test performed using active or inactive lever presses as dependent variables and Treatment as independent variable.

The gating protocol used for the classification of the different immune subpopulations and the criterion to estimate the percentage of positive cells for the different receptors studied are specified in Figure 1. Flow cytometry data were analysed with Student’s t-test for unrelated samples for each cell population and receptor. Cell number (total or receptor-expressing) was considered as the dependent variable and Treatment as the independent variable.

**Figure 1.**
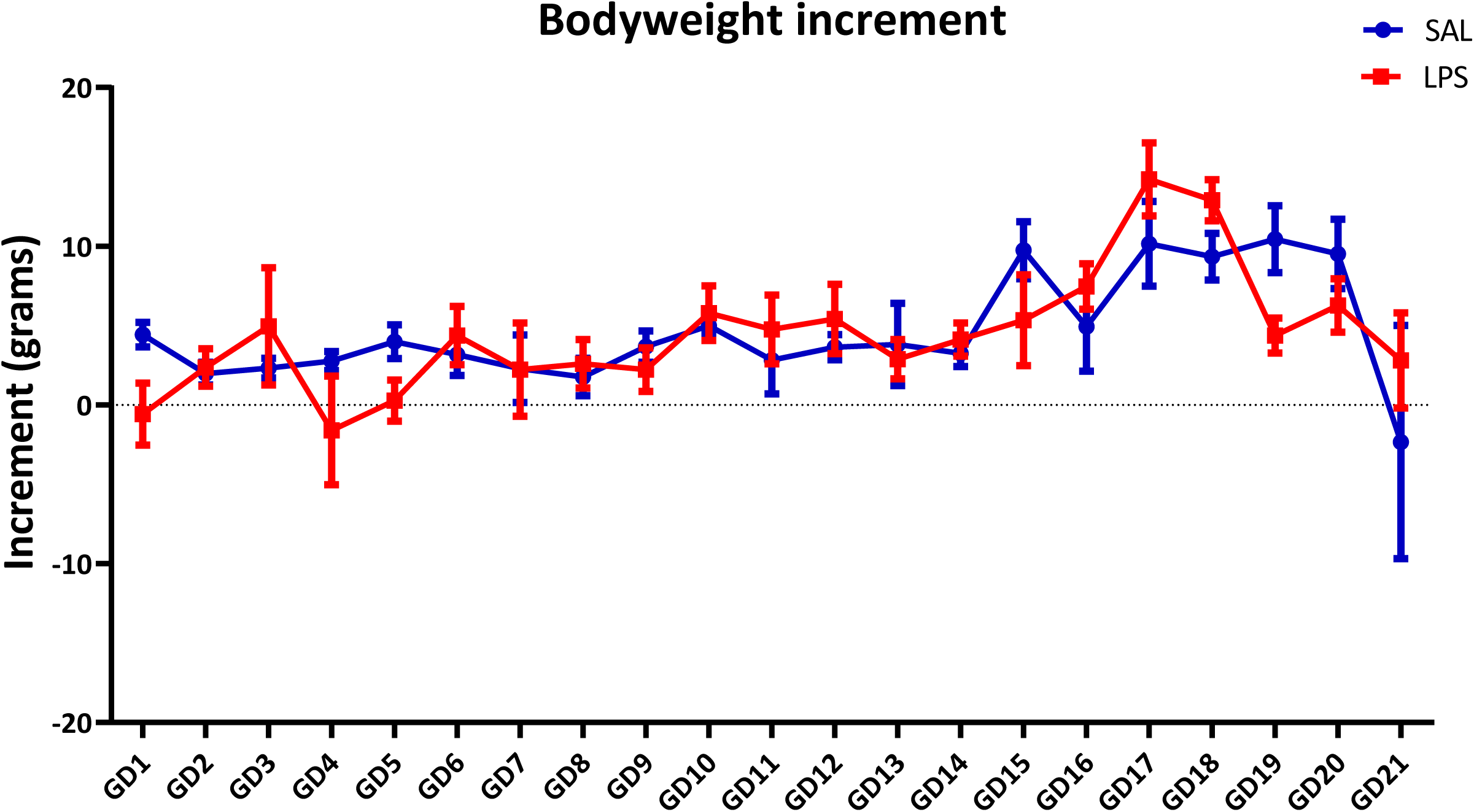
The body weight of the dams did not show any significant modification during pregnancy as a result of LPS exposure. No effects due to the Treatment or the Treatment * Gestational Day interaction were found.

Effect sizes are reported when appropriate. In the case of the Student’s t-tests, the effect size is estimated using Cohen’s d (the range of values being 0.2 = small effect, 0.5 = moderate effect, 0.8 = large effect: (Cohen, 1988). The partial η2 was used as an index of the effect size in ANOVAs (the range of values being 0.02 = small effect, 0.13 = moderate effect, 0.26 = large effect: (Miles and Shevlin, 2001)).

## Results

### 1. Daily LPS administration had no effects on the bodyweight of the dams

No significant differences were found among the treatments (F_(1,13)_=0.584, p=0.459) or due to the Treatment * Gestational Day interaction (F_(1,13)_=1.331, p=0.165) in the increment of body weight across gestation. However, the statistical analyses do show a significant effect for the Gestational day factor (F_(1,13)_ = 5.525, p< 0.0001; partial η2= 0.315) and a significant *Difference* contrast was found on GD 18 (p< 0.05, this contrast tests if there is a significant difference between one level of a variable and the previous one in a repeated-measures design). See Figure 2.

**Figure 2.**
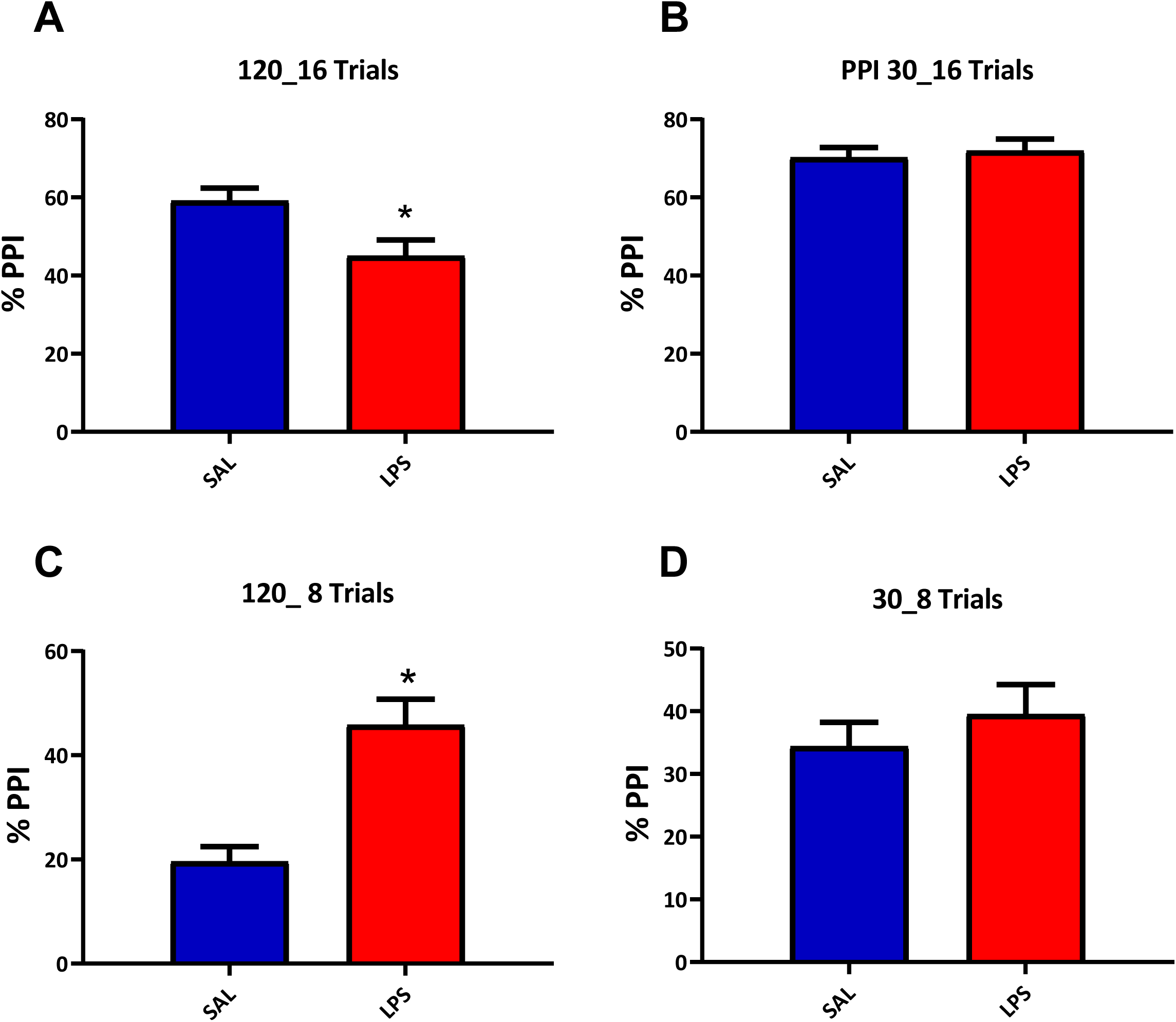
Percentage of prepulse inhibition in SAL or LPS-exposed offspring. There was a significant effect of the Treatment in the PPI120_16 (A) trials whereby LPS rats showed a weaker PPI than SAL controls. Conversely, in 120_8 (C) trials the opposite pattern emerges and LPS rats have a higher PPI index: * p< 0.05 compared to saline-controls. No significant effects of the Treatment were found in the other conditions tested.

### 2. LPS-exposed offspring showed a diminished PPI as compared to control rats in 120_16 trials

When tested at PD 120, LPS rats showed a significantly diminished PPI in PPI120_16 trials (t_60_=2.721, p< 0.05; Cohen’s d=0.069). Conversely, in the PPI120_8 trials, the opposite result was found whereby LPS-exposed rats displayed a higher PPI percentage (t_46.97_ =-4.528, p<0.001; Cohen’s d=0.116). See Figure 3 A-B.

**Figure 3.**
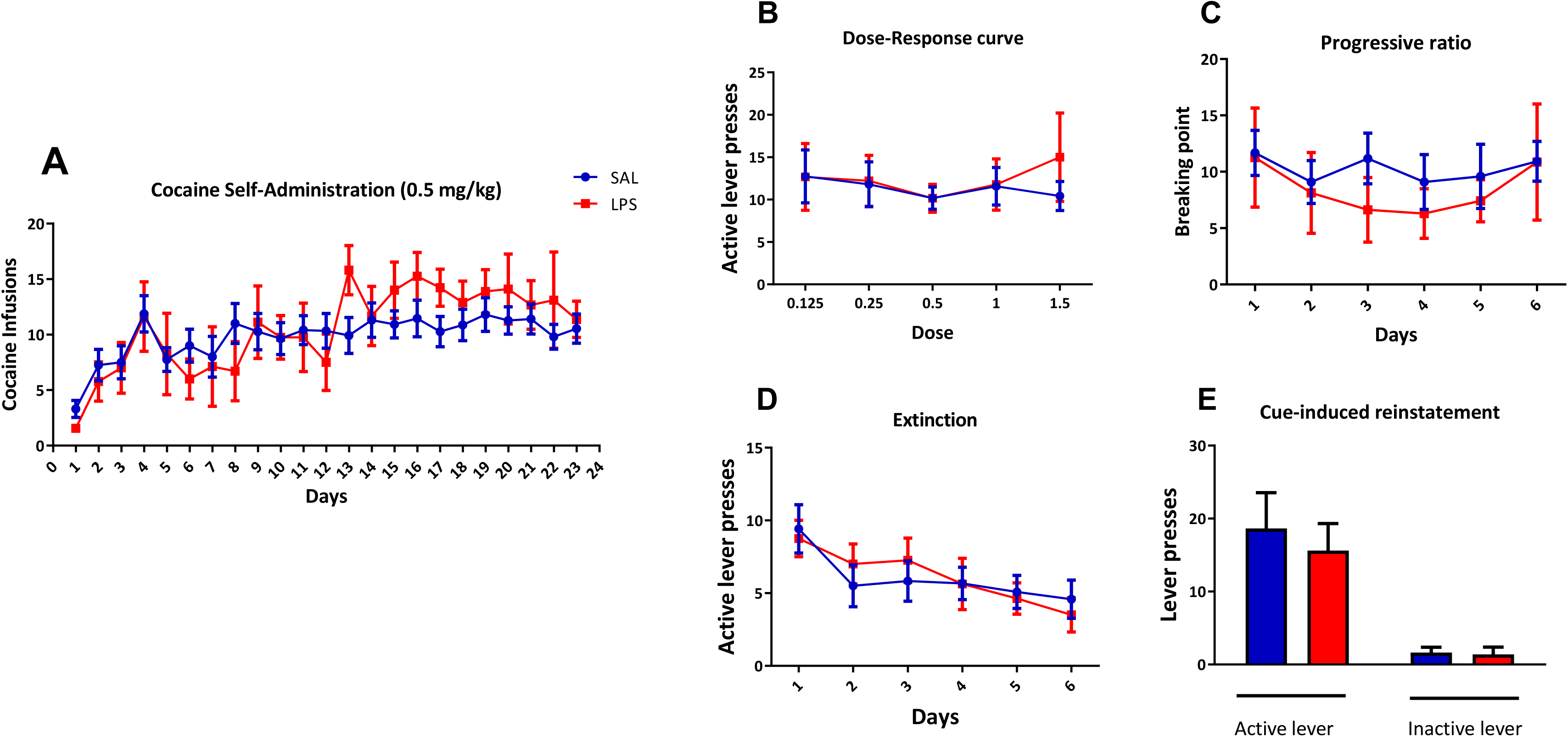
Cocaine self-administration. A) The figure shows the number of cocaine infusions earned during the acquisition phase under an FR1 schedule (0.5 mg/kg dose). Significant differences were found due to the Sessions factor, but no other differences were found in cocaine self-administration behaviour due to Treatment or the Session * Treatment interaction. B) Mean number of active lever presses per dose. There was no effect of Sessions, Treatment or their interaction. C) Breaking points in the progressive ratio schedule. No significant effects were found. D) Number of active lever presses during extinction. Animals progressively decreased their seeking behaviour over the extinction sessions, but no significant effects of Treatment or Sessions x Treatment interaction were observed. E) Number of active and inactive lever presses in the cue-induced reinstatement. No differences were found due to the Treatment.

No significant effects were found in 30_16 (t_61_ = −0.431, p= 0.668) or 30_8 trials (t_58_ = 0.845, p= 0.402) (Figure 3 C-D).

### 3. LPS-exposed offspring did not show increased intake of cocaine, motivation for its consumption or cue-induced drug seeking as compared to saline-exposed controls

Rats progressively increased their cocaine intake over sessions as suggested by the significant effect of the Sessions factor (F_(1,23)_=7.358, p<0.0001; partial η2=0.251). We found a significant *Difference* contrast from the eighth session onwards (p<0.05). However, there were no significant effects of the Treatment (F_(1,23)_=0.005, p=0.942) or the Treatment * Session interaction (F_(1,23)_=1.071, p=0.375) (Figure 4).

**Figure 4.**
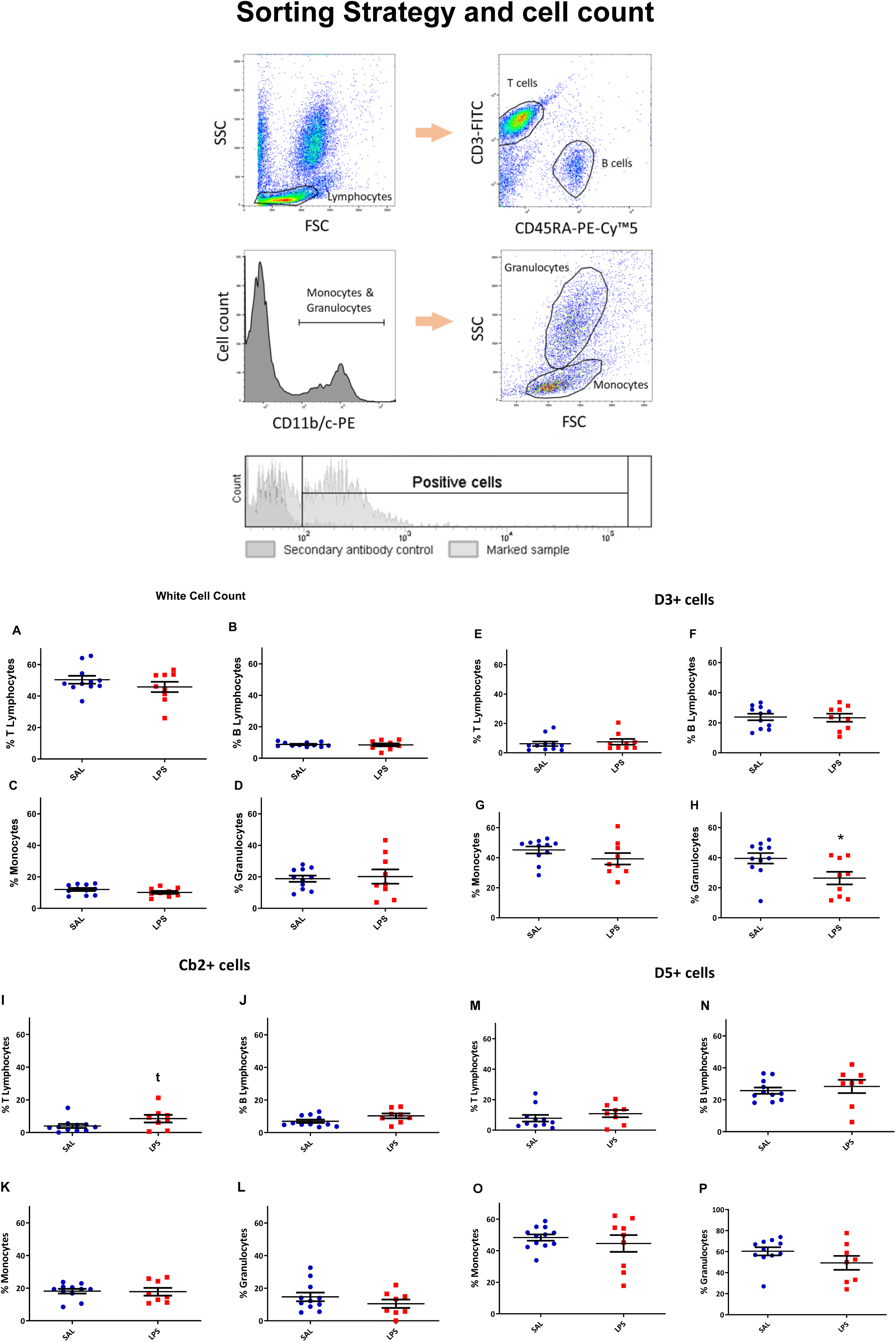
A) Classification of the immune cell subpopulations and characterisation of CB2, D3 and D5 positive cells. Lymphocytes were gated in the FSC vs SSC plot, then T cells were identified by the surface expression of CD3, while B cells were identified by the presence of CD45RA. Monocytes and granulocytes were first gated by selecting the CD11b/c positive cells and then were separated by their physical properties based on the FSC vs SSC plot. In order to quantify the CB2+, D3+ and D5+ cells among the gated subpopulations, we set a threshold based on the secondary antibody control. Then we subtracted the proportion of positive cells of the secondary antibody control from the positive cells of the marked sample. A-D) Percentage of immune cells in peripheral blood. E-F, I-J, M-N) Percentage of D3+, CB2+ and D5+ T and B lymphocytes, respectively, in SAL or LPS-exposed offspring and G-H, K-L and O-P) Percentage of D3+, CB2+ and D5+ monocytes and granulocytes. * Different from the saline group (p<0.05). t trend for a difference from the saline group (p=0.056).

In the dose response curve we did not find significant differences of the Sessions (F_(1,20)_ = 0.333, p= 0.855), Treatment (F_(1,20)_ = 0.019, p= 0.892) or their interaction (F_(1,20)_ = 0.673, p= 0.612) (Figure 5-A). No significant differences of the Sessions (F_(1,8)_ = 0.530, p= 0.703), Treatment (F_(1,18)_ = 1.940, p= 0.182) factors or the Treatment * Session interaction (F_(1,18)_ = 0.153, p= 0.954) (Figure 5-B) were found in the progressive ratio schedule.

During extinction rats decreased their lever pressing behaviour as highlighted by the significant effect of the Sessions factor (F_(1,19)_=6.774, p<0.0001; partial η2= 0.273). No effects were found due to the Treatment factor (F_(1,19)_=0.005, p=0.945) or Treatment * Session interaction (F_(1,19)_=0.674, p=0.596) (Figure 5-C).

There were no significant differences between the Treatments in active lever presses (t_14_=0.391, p=0.702) or inactive lever presses (t_13_=-0.457, p=0.655) during cue-induced reinstatement (Figure 5-D).

### 4. LPS-exposed offspring had fewer D3-expressing granulocytes than saline controls

First, we wanted to evaluate whether the schizophrenia-like phenotype produced by MIA had an effect on blood cell populations. The statistical analyses did not reveal differences between treatments in any studied population: B lymphocytes (t_19_ = 0.634, p= 0.534); T lymphocytes (t_19_ = −0.059, p= 0.953); monocytes (t_19_ = 1.072, p= 0.297) and granulocytes (t_11.368_ = −0.482, p= 0.639) (Figure 6 A-D).

With regard to the labelling performed for CB2 receptors in the different populations of immune cells, we did not find any significant differences between the treatments. However, there seems to be a trend for a Treatment effect in both types of lymphocytes. Analyses performed for CB2 receptors on T lymphocytes show a trend for up-regulation in LPS-exposed rats (t_18_ = −2.039, p= 0.056) as was also the case for B lymphocytes (t_18_=-1.982, p=0.063) (Figure 7 A-D).

We did not find any significant difference in monocytes (t_18_ = 0.368, p= 0.717) or granulocytes (t_18_ = 1.264, p= 0.223) (Figure 7 B-C).

We did find, however, a statistically significant lower number of D3 receptor-expressing granulocytes in the blood in LPS-exposed offspring (t_60_=2.721, p<0.05; Cohen’s d=1.121) (Figure 8-D). No differences were found between the treatments in the rest of populations for this receptor: T lymphocytes (t_19_ = 0.745, p= 0.465); B lymphocytes (t_19_ = 0.409, p= 0.687) or monocytes (t_19_ = 1.537, p= 0.141) (Figure 8 A-B-C).

Finally, no differences were found between treatments for D5 receptor-expressing cells in any of the populations studied: T lymphocytes (t_18_=-1.559, p=0.136); B lymphocytes (t_19_ =-0.801, p=0.433); monocytes (t_10.235_=0.665, p=0.521) or granulocytes (t_19_=1.657, p=0.114) (Figure 9 A-D).

## Discussion

In the experiments here reported we have used a widely used MIA model to recapitulates several aspects of the complex constellation of symptoms in schizophrenia to test if the motivation for cocaine, the sensitivity to the rewarding actions of the drug and the propensity to relapse were potentiated. We also explored the potential of some immune populations to serve as diagnostic tools or biomarkers of the disease. While there were no alterations in cocaine-related behaviours, consistent with our previous results (Santos-Toscano et al., 2016), we found a lower number of DR3D-expressing granulocytes and a trend for an increased number of CNR2-expressing lymphocytes in LPS-exposed rats.

Using the MIA approach, we have reproduced the typical reduction in PPI (in the PPI120_16 trials) observed in these models and patients. No alterations were detected in somatic growth during pregnancy, as evidenced by the weight gain of the dams. Patients with schizophrenia develop severe attention and sensory information processing deficits. PPI is an important example of the sensorimotor gating that occurs in the brain, and it is disrupted in individuals with schizophrenia, although such disruption can be reversed with atypical antipsychotics (Van den Buuse et al., 2003). As stated above, there is ample consensus in the literature that PPI is disrupted in MIA models of schizophrenia (Boksa, 2010; Fortier et al., 2007; Ozawa et al., 2006; Romero et al., 2010; Santos-Toscano et al., 2016; Wischhof et al., 2015). It is interesting to note that in the PPI120_8 trials, we have observed increased PPI in LPS-exposed rats. Increased PPI using short prepulse-pulse intervals has been documented before and calls for caution when using PPI as the exclusive index of the searched phenotype (Swerdlow et al., 2008).

There is clinical evidence for an increased risk of drug use among schizophrenic patients, mostly with nicotine, but also cocaine and other drugs (Khokhar et al., 2018); however, the causality of this relationship is not clear. There has been an increasing effort from preclinical researchers to provide some answers to this conundrum (Berg et al., 2014; Brady et al., 2008; Chambers and Self, 2002; Fletcher et al., 2018; Gallo et al., 2014; Karlsson et al., 2013; Ng et al., 2013; Ruda-Kucerova et al., 2017; Santos-Toscano et al., 2016; Sentir et al., 2018; Spano et al., 2010; Waterhouse et al., 2018) but most of the reports using self-administration methodologies have focused on pharmacological or neonatal ventral hippocampal lesions (NVHL). Arguably, MIA models better resemble some of the developmental factors affecting the proclivity to develop a disorder of the schizophrenic spectrum (Meyer, 2019), however, just a few MIA studies have focused on the voluntary consumption of drugs using self-administration methodologies (Liu et al., 2004; Santos-Toscano et al., 2016; Waterhouse et al., 2018). In a previous work, we showed that the acquisition of cocaine self-administration was unaltered in the offspring of LPS-exposed pregnant rats (Santos-Toscano et al., 2016). Here we wanted to expand these data by testing the motivation for cocaine using a progressive ratio schedule, the sensitivity to the unit dose of the drug and the resistance to extinction and proclivity to relapse using a cue-induced reinstatement procedure. In the present experiments, we also modified the LPS administration regime as compared to the one used by us in the past, to a more prolonged schedule (alternate days for the whole pregnancy) to test if this new MIA program resulted in more significant alterations.

Coincidentally with our previous report (Santos-Toscano et al., 2016), there was no effect of MIA on the acquisition of cocaine self-administration in the present experiments. Interestingly, we found no evidence for sensitivity to changes in the unit dose of cocaine in either group. This may have been derived from the prolonged acquisition program that we have used. Indeed, prolonged training schedules may result in the development of habit-like behaviour, which may not be sensitive to changes in the dose of the drug. Moreover, we have not found evidence for increased motivation for the drug using a progressive ratio schedule. There are conflicting data about the effects of MIA on the performance under progressive ratio schedules. For example, previous data using operant food self-administration also showed no alteration in food motivation in MIA LPS-exposed rats (Straley et al., 2017). However, when the immune activation took place at an earlier gestational day (GD12), the motivation for the food pellets under an amphetamine challenge was reduced in the absence of detectable disruptions in the dopaminergic system. Contrarily, the work by Millar and colleagues showed that Poly I:C administration at GD15 increased the motivation for food, possibly due to an inability to detect changes in the contingency between their behaviour and the resulting rewarding outcome (Millar et al., 2017). It must be noted that these experiments were performed using food pellets as reinforcers while we have examined the motivation for cocaine which may explain the differences in the results obtained. Lastly, whilst other authors (using the NVHL approach and cocaine) found increased response rates and delayed extinction-criterion attainment (Chambers and Self, 2002) we found no changes in extinction rates or in cue-induced reinstatement behaviour. Again, this divergence may be derived from the intrinsic differences in the developmental alterations that appear following a neonatal hippocampal lesion and those observed after MIA (Jones et al., 2011).

In summary, we have not observed any evidence for increased cocaine intake, motivation or relapse vulnerability in our MIA model of schizophrenia suggesting that the high rate of cocaine used among schizophrenia patients may be a result of other factors or mechanisms such as self-medication or increased stress from the environment.

The last experiments carried out in this study were aimed at identifying potential biomarkers for schizophrenia using the MIA model.

Many clinical studies have found differences in cytokine levels in schizophrenic patients and identified associations with the diagnosis of the disorder, the severity of the symptoms or the response to treatment (de Witte et al., 2014; Freudenreich et al., 2010; Loffler et al., 2010; Swiergiel and Dunn, 1999). In the light of the importance of the dopaminergic system in the etiopathogenesis of schizophrenia and the involvement of dopaminergic receptors in clinical biomarkers studies performed in schizophrenic patients (Buttarelli et al., 2011), we decided to focus on the different populations of immune cells sorted according to the expression of dopaminergic receptors, specifically D3 and D5. In addition, we also used the cannabinoid receptor type 2 (CNR2) surface expression as a means to find a population of immune cells that could be associated to the effects of MIA and could be used to identify susceptible individuals.

There were no differences produced by MIA in terms of the peripheral blood population numbers of the different cell populations measured here (B cells, T cells, granulocytes and monocytes). In contrast, clinical studies have shown that the populations of T lymphocytes and monocytes are increased in schizophrenic, chronic and acute patients, without observing differences in B lymphocytes (Cazzullo et al., 1998; Drexhage et al., 2011; Kelly et al., 2018). Therefore, it is conceivable that there is an overactivation of the immune system that is independent of the stages of the disorder. However, it remains to be clarified whether this overactivation is typical of the schizophrenic disorder or can be shared by other disorders since similar results have been found in patients with bipolar disorder (Sourlingas et al., 2003). The differences existing between our results and those of human studies may be related to the fact that we have used pan B and T cell markers while in the clinical setting a more detailed subset of B and T cells was studied.

As for the clinical studies that present results for dopamine receptors in immune cells, there seems to be more controversy. Several studies show an increase in the mRNA of the DRD3 in lymphocytes in schizophrenic patients (Boneberg et al., 2006; Ilani et al., 2001). However, this result may not be specific to schizophrenia as one study found that DRD3 mRNA levels were decreased among schizophrenic patient but also in patients with bipolar disorder (Vogel et al., 2004). Concerning DRD5, no differences in mRNA expression have been reported (Kwak et al., 2001). In our study, we found that the number of granulocytes but not lymphocytes expressing DRD3 was lower in LPS-exposed animals, with no alterations in DRD5-expressing leukocytes. It should be noted that dopaminergic receptors are not distributed in the same way in all immune cells. Indeed, the highest expression of these receptors is found in lymphocytes, with DRD3 and DRD5 being the most stable subtypes in the different populations (McKenna et al., 2002; Santambrogio et al., 1993). The specific effect in granulocytes that we observe may be related to the fact that LPS activates neutrophils (a granulocytic subpopulation). Bearing this in mind, future studies should determine which subpopulation of granulocytes are responsible for the drop in DRD3-expressing cells in MIA rats. Ideally, other models of schizophrenia which do not involve an immunogenic agent should be used to reproduce this result and rule out the possibility that the effect here reported may be directly linked to the immune actions of LPS and not the schizophrenic phenotype produced by MIA.

Finally, we found an upward trend in the CNR2 receptor-expressing lymphocytes. Interestingly, clinical studies have revealed that the expression levels of this receptor is diminished in peripheral blood mononuclear cells (PBMCs) (Bioque et al., 2013; Ferretjans et al., 2014). However, these results only appear in patients who had a previous history of cannabis use or who were in treatment. Another study where the participants reported no abuse/dependence of marihuana, the levels of CNR1 and CNR2 mRNA were significantly higher in participants with schizophrenia, however, this study did not distinguish among the different populations of PBMCs (Chase et al., 2016). As regards this, it is crucial to study the expression of the cannabinoid receptors in PBMCs of patients who have not been exposed to any treatment with antipsychotics.

## Conclusions

In a widely used MIA model of schizophrenia there is no evidence for increased cocaine self-administration, motivation for the drug, sensitivity to the dose, extinction or cue-induced reinstatement of drug-seeking behaviour, suggesting that the clinical relationship between increased drug use and schizophrenia does not originate as a result of the primary neural alterations associated to the disease. In considering the results obtained in our flow cytometry experiments, we suggest DRD3+granulocytes and CNR2+ lymphocytes as potential biomarkers in schizophrenia that should be further validated.

## Contributors

RS-T, MU and EB performed the experiments. RS-T, MU also performed the statistical analyses. RS-T wrote the first draft of the manuscript. RS-T, MU and AH-M wrote the final version of the manuscript. EA and AH-M designed the experiments, provided funding and supervised the final version of the manuscript. All authors have approved the final manuscript.

## Acknowledgements

We thank Rosa Ferrado, Luis Carrillo, Gonzalo Moreno and Alberto Marcos for their excellent technical assistance.

